## Supplementary Information for "Effects of cryo-EM cooling on structural ensembles"

#### Supplementary Methods

##### Rmsf decrease in different parts of the ribosome-complex

To measure the narrowing of structural ensembles during cooling, we used the quantiles of the rmsf distributions (Figs. 2, 3). However, when using rmsf distributions obtained from the whole set of atoms, the information of where the atoms are located and the information to which kind of molecule the atoms belong to are lost. To address this limitation, we divided the ribosome-complex atoms into a set containing only RNA atoms, a set of protein atoms and separately calculated the rmsf distributions before and after the longest T-quench simulations ($\tau_c$=128 ns). Further, we divided the set of atoms into five shells around the center of geometry of the ribosome, where each set contains the same number of atoms. For each of these sets, we calculated rmsf distributions before and after the 128-ns T-quench simulations (Fig. S2a). Figure S2b shows the rmsf difference before and after cooling. Mean values and standard deviations were obtained as described for the rmsf distributions containing all atoms (Methods).

##### Protein backbone and side-chain conformations

To check if the backbone and side-chain conformations of the proteins in the ribosome complex are affected by the cooling rate, we extracted backbone dihedral angles $\Phi$ and $\Psi$ and side-chain rotamer angles $\chi_1$ and $\chi_2$ from the ensembles before, during, and at the end of all the T-quench simulations. To that end, we applied the program *gmx chi* of the gromacs package[1] to the ensembles of structures described in the Methods section *Structural heterogeneity during T-quench simulations*. For each ensemble of structures, the $\Phi$ and $\Psi$ and angles from all protein residues in all structures were sorted into 2-d bins (180 for each angle). Ramachandran plots of the ensemble before cooling and at the end of the 128-ns T-quench simulation are shown in Fig. S4a. To quantify the narrowing of the protein backbone conformations, we calculated Shannon entropies of the angle distributions. To measure the statistical uncertainty, we used bootstrapping, as described for the structural heterogeneity, and calculated the entropy for each ensemble. The mean values and standard deviations are shown in Fig. S4b. For amino acids containing $\chi_1$ and $\chi_2$ rotamers, the Shannon entropies were calculated as described for the backbone dihedral angles. Figure S4c show the entropy change with respect to the ensemble of structures prior to cooling.

#### Numerical solution of the heat equation for a water layer

To solve the heat equation, we used the Crank-Nicolson method (central difference in time and in space, CTCS)[2] with discretized time and space $T(i\Delta x, n\Delta t) = T_i^n$, with $\Delta x = 1$ nm and $\Delta t = 0.01$ ns:

$$\frac{T_i^{n+1} - T_i^n}{\Delta t} = \frac{1}{2}\Bigg( \frac{[\alpha(x)\partial_x T(x,t)]_{x=(i+\frac{1}{2})\Delta x}^{t=(n+1)\Delta t} - [\alpha(x)\partial_x T(x,t)]_{x=(i-\frac{1}{2})\Delta x}^{t=(n+1)\Delta t}}{\Delta x} + \frac{[\alpha(x)\partial_x T(x,t)]_{x=(i+\frac{1}{2})\Delta x}^{t=n\Delta t} - [\alpha(x)\partial_x T(x,t)]_{x=(i-\frac{1}{2})\Delta x}^{t=n\Delta t}}{\Delta x} \Bigg).$$

With $l_i = \frac{\Delta t}{4\Delta x^2}\alpha\left((i-\frac{1}{2})\Delta x\right)$ and $r_i = \frac{\Delta t}{4\Delta x^2}\alpha\left((i+\frac{1}{2})\Delta x\right)$, the discretized heat equation can be written as

$$-l_i T_{i-1}^{n+1} + (1 + r_i + l_i) T_i^{n+1} - r_i T_{i+1}^{n+1} = l_i T_{i-1}^n + (1 - r_i - l_i) T_i^n + r_i T_{i+1}^n.$$

For vectors $\boldsymbol{T}^{n+1} = (T_2^{n+1}, \cdots, T_{N-1}^{n+1})^\intercal$ and $\boldsymbol{T}^n = (T_2^n, \cdots, T_{N-1}^n)^\intercal$, this equation can be written in matrix form $A \cdot \boldsymbol{T}^{n+1} = B \cdot \boldsymbol{T}^n + \boldsymbol{b}$, where $A$ and $B$ are $(N-2) \times (N-2)$ matrices:

$$A = \begin{pmatrix} 1+r_2+l_2 & -r_2 & 0 & \cdots & & & & 0 \\ -l_3 & 1+r_3+l_3 & -r_3 & 0 & \cdots & & & 0 \\ 0 & -l_4 & 1+r_4+l_4 & -r_4 & 0 & \cdots & & 0 \\ \vdots & & & & \ddots & & & \vdots \\ & & & & & -l_{N-2} & 1+r_{N-2}+l_{N-2} & -r_{N-2} \\ 0 & \cdots & & & & 0 & -l_{N-1} & 1+r_{N-1}+l_{N-1} \end{pmatrix},$$

$$B = \begin{pmatrix} 1-r_2-l_2 & r_2 & 0 & \cdots & & & & 0 \\ l_3 & 1-r_3-l_3 & r_3 & 0 & \cdots & & & 0 \\ 0 & l_4 & 1-r_4-l_4 & r_4 & 0 & \cdots & & 0 \\ \vdots & & & & \ddots & & & \vdots \\ & & & & & l_{N-2} & 1-r_{N-2}-l_{N-2} & r_{N-2} \\ 0 & \cdots & & & & 0 & l_{N-1} & 1-r_{N-1}-l_{N-1} \end{pmatrix}.$$

The $N-2$ dimensional vector $\boldsymbol{b}$ contains the temperature $T_b = 90$ K at the boundary grid points $i = 1$ and $i = N$, $\boldsymbol{b} = \frac{T_b \Delta t}{2\Delta x^2}(\alpha(\Delta x), 0, \ldots, 0, \alpha(N\Delta x))^\intercal$. After multiplying with the inverse of matrix $A$, we get

$$\boldsymbol{T}^{n+1} = A^{-1}B \cdot \boldsymbol{T}^n + A^{-1}\boldsymbol{b}.$$

The starting temperature profile $\boldsymbol{T}^0$ was set to a temperature of 90 K for the ethane layer and to 277.15 K for the water layer (Fig. 1a, blue line). Next, equation  was iterated until the temperature in the center of the profile dropped below 90.1 K (Fig. 1c).

#### Metropolis sampling with Bayesian inference

The models of the cooling process model1, model2, and model3 have different sets of parameters $\boldsymbol{p}$. Here, the aim is to obtain the probability distribution $P(\boldsymbol{p} \mid \boldsymbol{Q}, \boldsymbol{S})$ of the parameter set $\boldsymbol{p}$ given the rmsf quantiles obtained from the T-quench simulations $\boldsymbol{Q} = \{\boldsymbol{Q}_{\tau_c} \mid \tau_c \in \boldsymbol{\tau}_c\}$, where $\boldsymbol{\tau}_c$ is the set of cooling time spans used for training the model, e.g., $\boldsymbol{\tau}_c = [0.1\text{ns}, \ldots, 128\text{ns}]$. The standard deviations of the rmsf values are given by $\boldsymbol{S} = \{\boldsymbol{S}_{\tau_c} \mid \tau_c \in \boldsymbol{\tau}_c\}$ as described above. According to Bayes' theorem,

$$P(\boldsymbol{p} \mid \boldsymbol{Q}, \boldsymbol{S}) \propto P(\boldsymbol{Q}, \boldsymbol{S} \mid \boldsymbol{p}) \cdot P(\boldsymbol{p}), \tag{1}$$

with the likelihood $P(\boldsymbol{Q}, \boldsymbol{S} \mid \boldsymbol{p})$ and the prior probability of the parameters $P(\boldsymbol{p})$. The likelihood is given by

$$P(\boldsymbol{Q}, \boldsymbol{S} \mid \boldsymbol{p}) = \prod_{i=1}^{5} \prod_{\tau_c \in \boldsymbol{\tau}_c} P(\mathbf{Q}_{\tau_c,i}, \mathbf{S}_{\tau_c,i} \mid \boldsymbol{p}),$$

where $\mathbf{Q}_{\tau_c,i}$ and $\mathbf{S}_{\tau_c,i}$ are the $i$-th rows of $\mathbf{Q}_{\tau_c}$ and $\mathbf{S}_{\tau_c}$, respectively. For each quantile $i$ and each cooling time span $\tau_c$, the likelihood is given by

$$P(\mathbf{Q}_{\tau_c,i}, \mathbf{S}_{\tau_c,i} \mid \boldsymbol{p}) = \prod_{j=1}^{11} \frac{1}{\sigma_{i,\tau_c}(t_j)\sqrt{2\pi}} \exp\left(-\frac{(Q_{i,\tau_c}(t_j) - \mathrm{rmsf}_{\mathrm{model}}(\boldsymbol{p}))^2}{\sigma^2_{i,\tau_c}(t_j)}\right),$$

where $\mathrm{rmsf}_{\mathrm{model}}$ is taken from equation 5 (Methods), equation 7 (Methods), and equation 8 (Methods) for model1, model2, and model3, respectively.

For the prior distributions of the parameters, we used uniform distributions. For model1 and model3, uniform distributions between 0 $\mathrm{MJ/mol/nm^2}$ and 30 $\mathrm{MJ/mol/nm^2}$ and between between 0 Å and 10 Å were used for parameters $c$ and $d$, respectively. For model2 and model3, uniform distributions between 0 Å and 10 Å, between 0 kJ/mol and 10 kJ/mol were used for parameters $\Delta x$ and $\Delta G^{\ddagger}$. For $\Delta G$ (model2 and model3), we used a uniform distribution between 0 kJ/mol and $-k_b T_h \log(1/0.99 - 1)) \approx 10.6$ kJ/mol such that the occupancy of state A is at least 1% at $T_h$ before cooling.

The probability distribution $P(\boldsymbol{p} \mid \boldsymbol{Q}, \boldsymbol{S})$ was sampled using the Metropolis Monte Carlo algorithm[3] with $10^6$ steps. To that aim, we used the function $f(\boldsymbol{p}) = P(\boldsymbol{Q}, \boldsymbol{S} \mid \boldsymbol{p}) \cdot P(\boldsymbol{p})$ which is proportional to the probability distribution (compare equation 1).

Initial values of parameters $d$ and $\Delta x$ were set to 1 Å for model variants which have one value for all quantiles and to 1 Å, 2 Å, 3 Å, 4 Å, and 5 Å for variants which have one value per quantile. Initial values of parameters $c$ were set to 5 $\mathrm{MJ/mol/nm^2}$ and initial values of parameters $\Delta G$ and $\Delta G^{\ddagger}$ were set to 5 kJ/mol.

For each metropolis step, $N_p$ substeps were carried out, where $N_p$ is the number of free parameters of the model variant. In each substep, a new value for one of the parameters was drawn from a normal distribution centered on the current value with a standard deviation of $\sigma_p$. For parameter $c$, $\sigma_p$ was set to 1 $\mathrm{MJ/mol/nm^2}$. For $d$, $\sigma_p$ was set to $2 \cdot 10^{-5}$ Å for variants with one value of $d$ for all quantiles and to $2 \cdot 10^{-5}$ Å, $4 \cdot 10^{-5}$ Å, $6 \cdot 10^{-5}$ Å $8 \cdot 10^{-5}$ Å, and $10 \cdot 10^{-5}$ Å for variants with one value of $d$ for each quantile. For $\Delta x$, $\sigma_p$ was set to $1 \cdot 10^{-4}$ Å for variants with one value of $\Delta x$ for all quantiles and to $1 \cdot 10^{-4}$ Å, $2 \cdot 10^{-4}$ Å, $3 \cdot 10^{-4}$ Å $4 \cdot 10^{-4}$ Å, and $5 \cdot 10^{-4}$ Å for variants with one value of $\Delta x$ for each quantile. For model2, $\sigma_p$ was set to 0.05 kJ/mol and 1 kJ/mol for parameters $\Delta G$ and $\Delta G^{\ddagger}$, respectively. For model3, $\sigma_p$ was set to 0.2 kJ/mol and 2 kJ/mol for parameters $\Delta G$ and $\Delta G^{\ddagger}$, respectively. In each substep, the function $f$ was evaluated with the new parameter, and the ratio $\alpha$ of the new and previous value of $f$ was used as the acceptance ratio: If $\alpha > 1$, the new parameter was accepted. If $\alpha < 1$, a random number $u$ between 0 and 1 was drawn and the new parameter was accepted if $u \leq \alpha$ and rejected otherwise.

#### Finding optimal model variants

The three models of the cooling process have different sets of parameters. When we use all model parameters for each quantile separately, model1, model2, and model3 have 10, 15, and 25 free parameters, respectively. For different variants of the models, we set different parameters to be the same for all quantiles. To test how well a model variant reproduces and predicts the rmsf values obtained from the T-quench simulations, we used a cross-validation approach. For each model variant, first, we applied Metropolis sampling with Bayesian inference (as described above) where the likelihood function only depends on the rmsf values from the T-quench simulations with cooling time spans between 0.1 ns and 64 ns. Next, after omitting the first 20% of the Metropolis steps, 1000 steps were randomly chosen, the values of the parameters were extracted and 1000 rmsf curves were calculated from the model for all cooling time spans. To check how well the model variant reproduces the rmsf values used for training, root mean square deviations (rmsds) of model rmsf values from those obtained from the T-quench simulations were calculated for cooling time spans 0.1–64 ns (Fig. S5, blue). To check how well the model variant predicts rmsf values not included in the training, rmsds of model rmsf values from T-quench simulation values were calculated for the cooling time span 128 ns (Fig. S5, red).

From the tested model variants, we chose the one with the lowest cross-validation rmsd for further analysis. For model1 and model2, we tested all possible variants (Fig. S5a,b) and we chose the model1 variant with five parameters for $d$ and one for $c$ and the model2 variant with five parameters for $\Delta x$ and $\Delta G^{\ddagger}$ and one for $\Delta G$. For model3, several variants showed similar cross-validation rmsds and among these variants we chose the one with the fewest free parameters, namely five parameters for $d$ and one for $c$, $\Delta x$, $\Delta G^{\ddagger}$, and $\Delta G$.

#### Comparison of models

To check how well the model recreates and predicts the rmsf values obtained from the T-quench simulations, we trained the model on different sets of cooling time spans (Fig. 3e). Next for each set of cooling time spans, after omitting the first 20% of steps, from 1000 randomly chosen Metropolis steps, the values of the parameters were extracted and 1000 sets of rmsf curves for all cooling time spans $\tau_c$ were calculated from the model. Then, rmsds of the rmsf values obtained from the model from those obtained from the T-quench simulations were calculated for all cooling time spans (Fig. 3e). The rmsf curves obtained from the models trained on all cooling time spans are shown in Fig. 3d (colored lines). Probability densities of the parameters obtained from two independent calculations are shown in Fig. S6a.

### Supplementary Results

#### Effect of additional sampling on rmsf during T-quench simulations

The T-quench simulations were started from 41 structures extracted from the 277.15-K simulation at intervals of 50 ns (1000 ns, 1050 ns,..., 3000 ns). During the T-quench simulations, we observed a decrease in rmsf values. However, this decrease might be partially canceled by the small rmsf increase due to additional sampling as observed in the 277.15-K simulations (compare time intervals 1-3 $\mu$s and 1.1-3.1 $\mu$s in Fig. S1b). For each cooling time span $\tau_c$ and for each time point $t_j$ of all the 41 T-quench trajectories, we extracted the 41 structures and calculated the rmsf distribution (Methods, Fig. 2c). Here, we additionally, calculated the rmsf change occurring during the additional sampling at T=277.15 K, by extracting a set of 41 structures from the 277.15-K trajectory at shifted times ($t_j$+1000 ns, $t_j$+1050 ns,..., $t_j$+3000 ns). This rmsf change we then subtracted from the rmsf values obtained from the T-quench simulations (Fig. S8). As expected, the resulting rmsf values (red) are generally lower than the rmsf values obtained from the T-quench simulations (black), especially for longer T-quench simulations (e.g., $\tau_c$=128 ns). However, because of the decreasing temperature in the T-quench simulations, the rmsf increase due to the additional sampling is expected to be smaller in the T-quench simulations compared to the 277K simulations. Therefore, the rmsf values obtained from the subtraction (red) represent an upper limit of the rmsf decrease during cooling. Since, we do not know where in the range between the rmsf values obtained from the T-quench simulations (black) and the rmsf values obtained from the subtraction (red), we used the T-quench values for our further analysis to make sure that we do not overestimate the effect of cooling.

#### Pre-exponential factor for model2 and model3

For model2 and model3, a temperature-dependent pre-exponential factor $\kappa\left(\frac{T}{T_h}\right)^{\nu}$ was used, where $T$ is the current temperature and $T_h$ is the temperature prior to cooling. For the temperature exponent $\nu$, we tested values 0.5, 1, and 2. For each value of $\nu$, we obtained the model parameters with Metropolis sampling and calculated the deviation between rmsf obtained from the MD simulation results and the rmsf values obtained from the model. The choice of $\nu$ did not affect how well the kinetic models agree with the MD simulation data (Fig. 3e) and we used $\nu$ =1 for the further analysis.

For model3, we tested values of $\kappa$ between 0.01 ns$^{-1}$ and 10 ns$^{-1}$, obtained the model parameters with Metropolis sampling and calculated the deviation between rmsf obtained from the MD simulation results and the rmsf values obtained from the model (Fig. S9). The deviation decreases with increasing $\kappa$ values and stays constant after reaching 1 ns$^{-1}$, suggesting a lower limit $\kappa \geq$1 ns$^{-1}$. Since larger values of $\kappa$ result in larger barrier height $\Delta G^{\ddagger}$ our results using a $\kappa$ value of 1 ns$^{-1}$ provide a lower limit of $\Delta G^{\ddagger}$.

#### Effect of cooling on conformational modes

To address the question if the distribution of conformations along dominant conformational modes is affected by cooling, we first obtained conformational modes using Principal Component Analysis (PCA)[4]. To that aim, we extracted structures at 5 ps intervals from the trajectory at T=271.15 K (1-3µs). To calculate the covariance matrix, we used the coordinates of P and CA atoms of RNA strands and proteins, respectively. The eigenvalue of each eigenvector of the covariance matrix corresponds to the variance of the projections onto this eigenvector, such that eigenvectors with large eigenvalues describe dominant conformational modes of motion. To restrict the analysis to conformational modes which contribute markedly to the overall motion, we chose the 212 eigenvectors whose eigenvalues are larger than 1 Å for further analysis. Next, for each of these eigenvectors, we projected all structures from which the T-quench simulations were started onto the eigenvector and calculated the standard deviation of the projections $\sigma_{\text{before}}$. For each cooling time span $\tau_c$, we then also projected the structures at the end of the T-quench simulations and calculated the standard deviation $\sigma_{\text{after}}$. The normalized difference between the standard deviations $\Delta\sigma = (\sigma_{\text{after}} - \sigma_{\text{before}})/\sigma_{\text{before}}$ for the eigenvectors as a function of their eigenvalues is shown in Fig. S10a.

For rapid conformational changes and given sufficient statistics, we would expect that the distributions of projections become narrower during cooling ($\Delta\sigma < 0$) and that this effect is more pronounced for longer cooling time spans. Figure S10b shows the percentage of eigenvectors for which $\Delta\sigma$ is below zero. We do not see a clear trend of narrowing, which could be explained by the conformational changes being too slow or a consequence of limited statistics. This result suggests that the local changes measured by the rmsf, where we see an effect, converge faster than the correlated motions involving the whole ribosome.

### Supplementary Figures

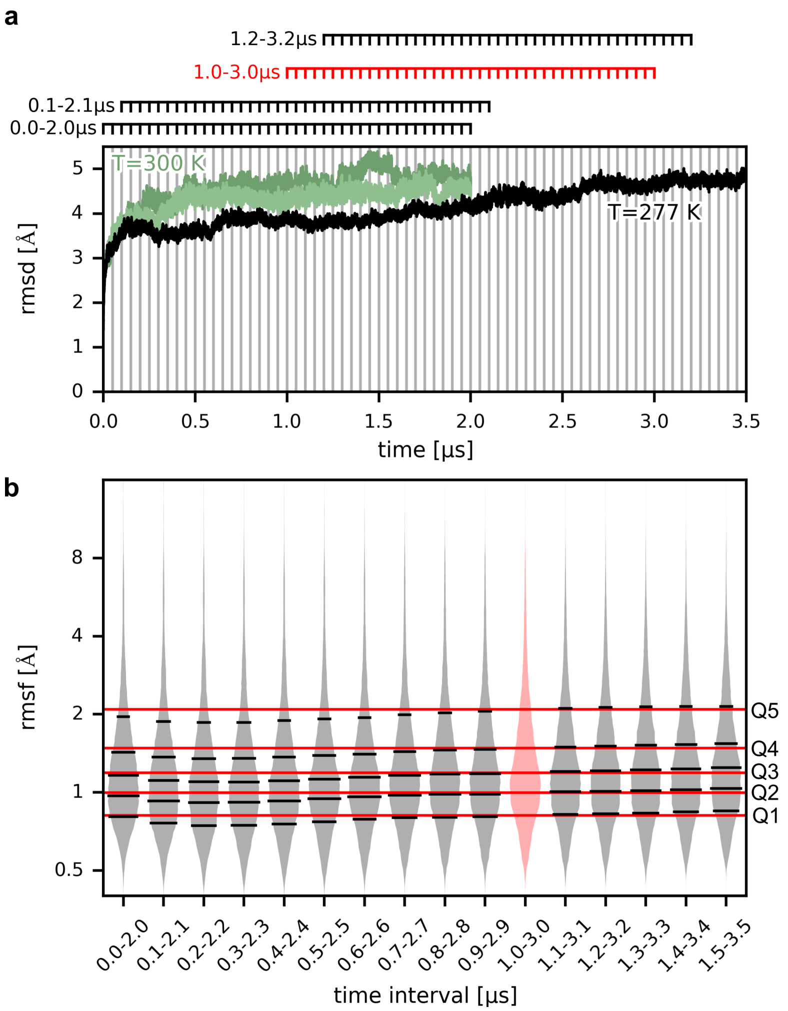


Figure S1: **Equilibration of the ribosome·EF-Tu complex at 277.15 K prior to cooling.** (**a**) Root mean square deviation (rmsd) as a function of simulation time (black line). Rmsd of two simulations of the same system at T=300 K (green lines) described earlier[5]. Time points of extracted snapshots are indicated by vertical lines (every 50 ns). Horizontal lines (top) indicate time intervals of 2 $\mu$s length. (**b**) The rmsf values of the ribosome·EF-Tu complex for ensembles of 41 structures obtained from different time intervals. Histograms of the rmsf values are shown as areas (grey, light red). The highlighted ensemble (red) was used for T-quench simulations. Horizontal lines (black, red) indicate the positions of the 6-quantiles (Q1–Q5) of each rmsf distribution.

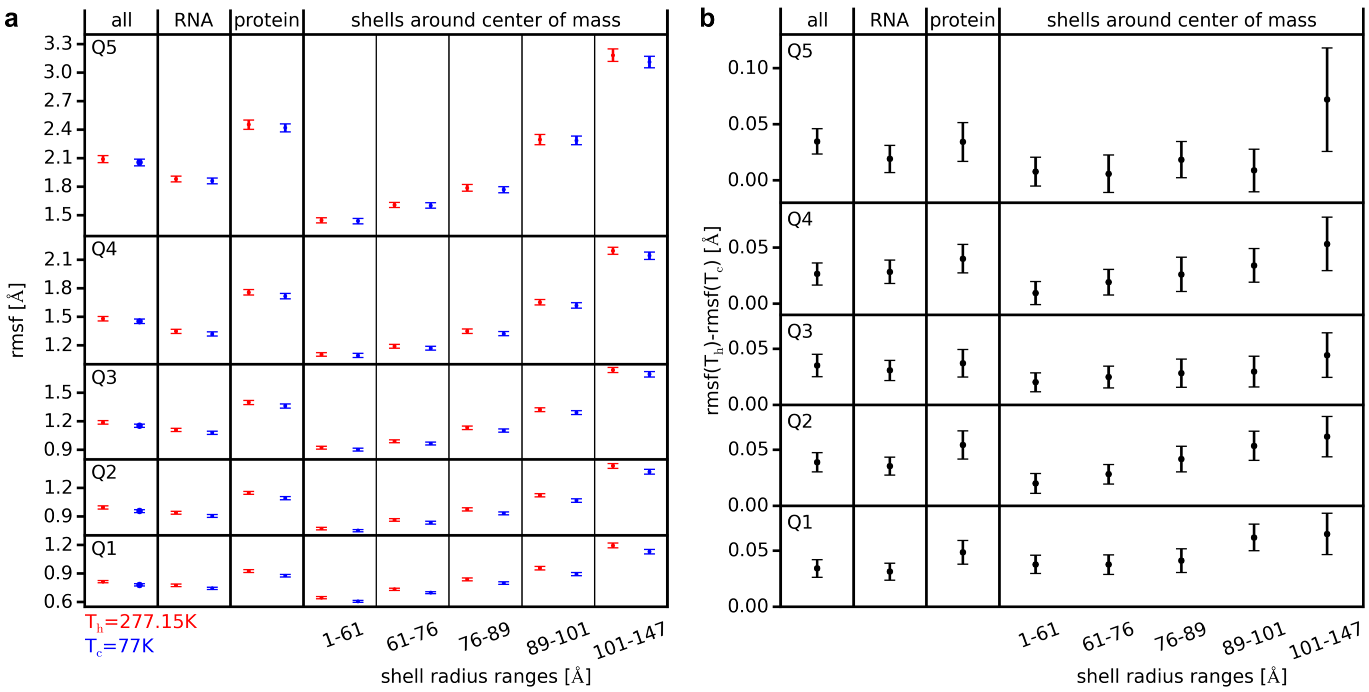


Figure S2: **Rmsf decrease in different parts of the ribosome·EF-Tu complex.** (**a**) Quantiles of the rmsf distribution before cooling (red) and after cooling (blue) obtained from the 128-ns T-quench simulations are shown for different sets of heavy atoms: all atoms, RNA atoms (rRNA+mRNA+tRNA), protein atoms (ribosomal proteins+EF-Tu), and five shells around the center of geometry of the ribosome complex. Mean values and standard deviations obtained from bootstrapping are depicted as circles and vertical bars, respectively. (**b**) Differences between the quantiles before and after cooling in (a).

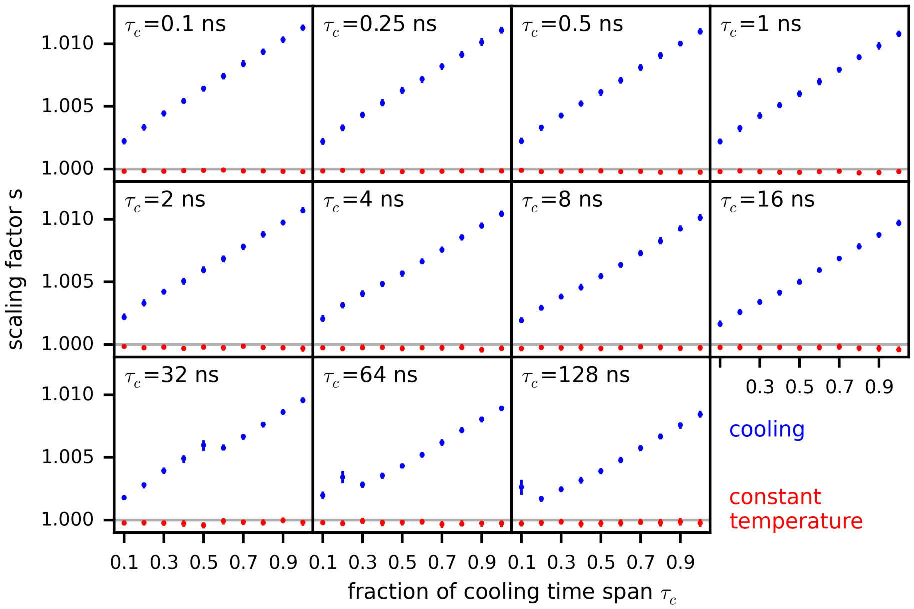


Figure S3: **Volume reduction during cooling.** For each cooling time span and each time point of the cooling trajectories, the scaling factor $s$ which minimized the average rmsd between the scaled conformations and the starting conformations of the respective simulations is shown (mean: blue circles, standard deviation: blue vertical lines). As a control, the same analysis was carried out for conformations extracted from the 277.15-K simulation (red circles and lines).

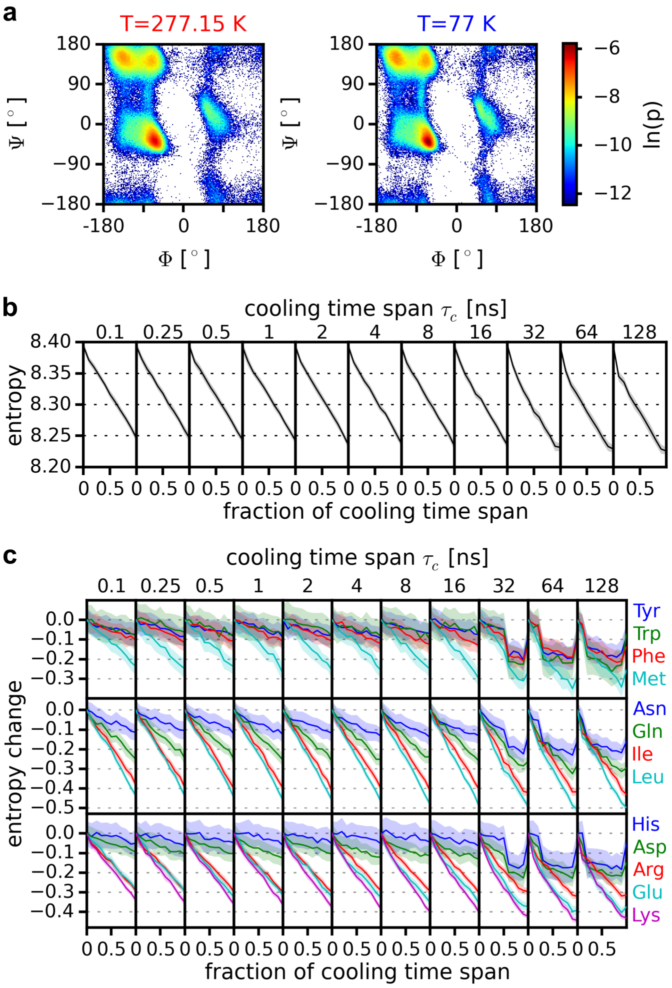


Figure S4: **Protein backbone and side-chain conformations during cooling.** (**a**) Ramachandran plot of protein backbone dihedral angles Φ and Ψ obtained from structural ensembles at T=277.15 K and and at T=77 K after T-quench simulations with a cooling time span $\tau_c$= 128 ns. (**b**) Shannon entropy of the distribution of Φ and Ψ angles during the T-quench simulations with different cooling time spans $\tau_c$. Mean values and standard deviations obtained from bootstrapping are shown as black lines and grey areas, respectively. (**c**) Shannon entropy change of the distribution of side-chain rotamer angles $\chi_1$ and $\chi_2$ during the T-quench simulations with different cooling time spans $\tau_c$. For each amino acid, mean values and standard deviations obtained from bootstrapping are shown as lines and areas, respectively.

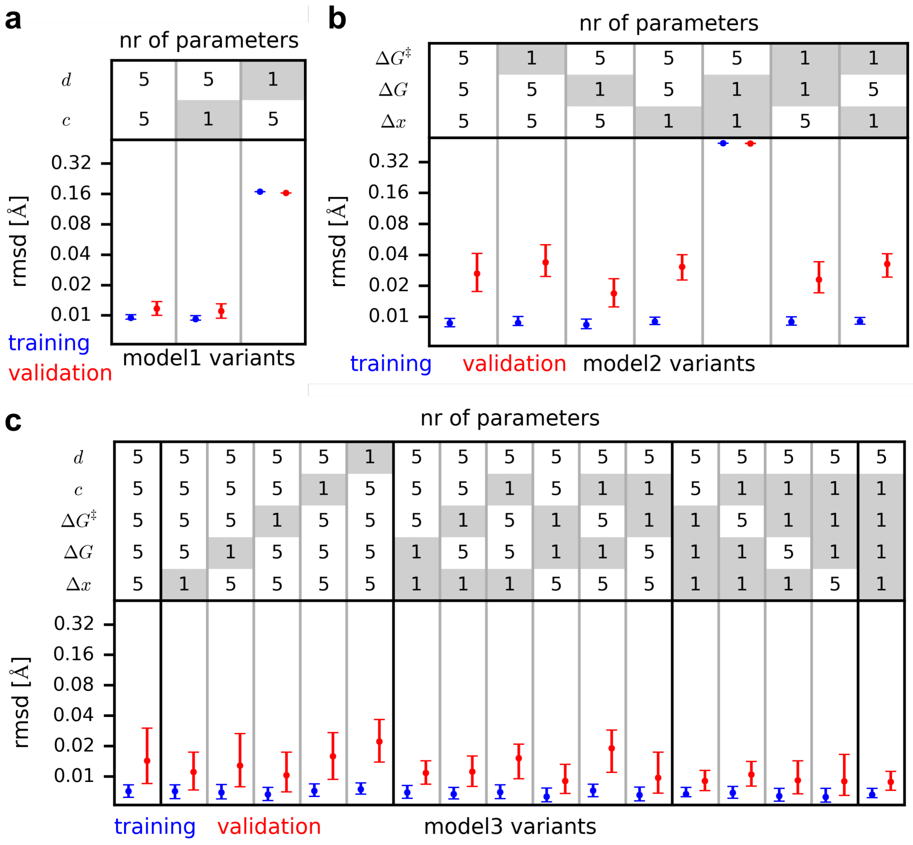


Figure S5: **Model parameter optimization.** Different variants of model1 (**a**), model2 (**b**), and model3 (**c**) with different numbers of the parameters $\Delta x$, $\Delta G$, $\Delta G^{\ddagger}$, $c$, and $d$ were trained on the rmsf values for cooling time spans between 0.1 ns and 64 ns (Fig. 2c, black lines). If the number of parameters is 5, there is one parameter for each quantile. If the number is 1, there is one parameter for all quantiles. The rmsds between the rmsf values calculated from the model and the training set are shown in blue (bars indicate the 95 % confidence interval). For validation, the rmsds between the rmsf values predicted by the model for cooling time span 128 ns and the rmsf values from the respective MD simulations are shown in red.

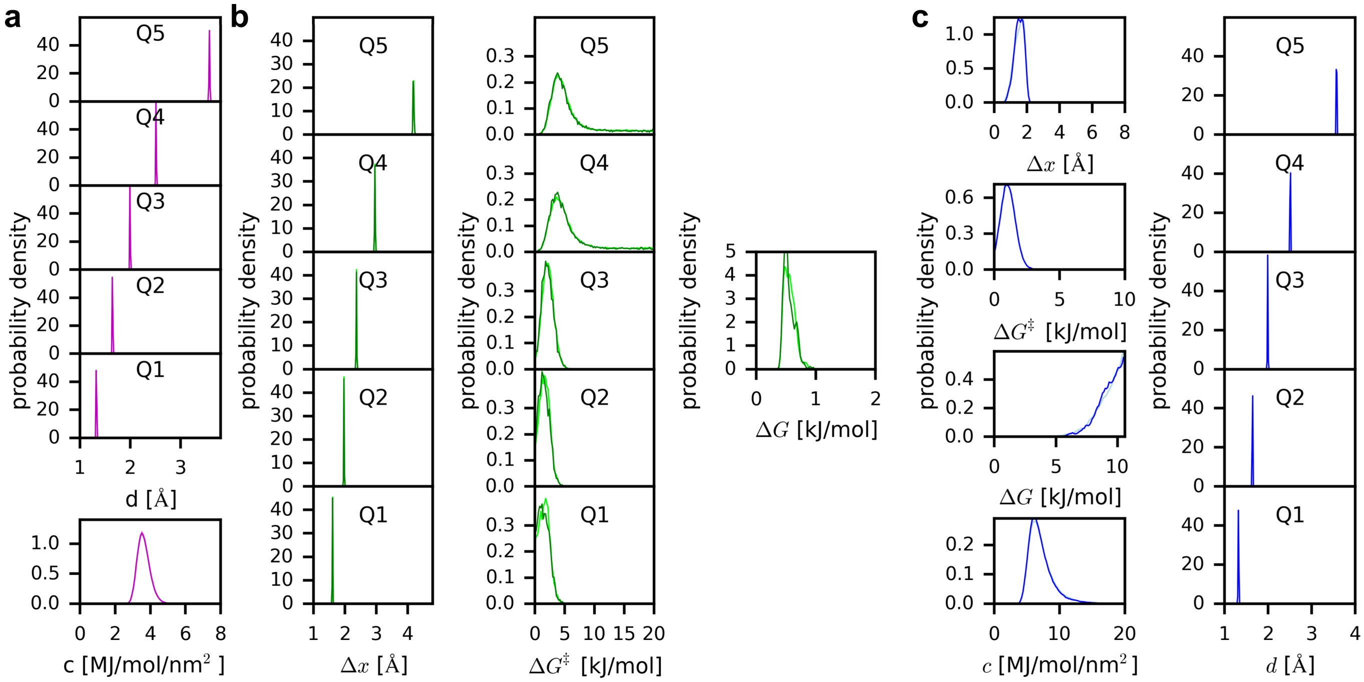


Figure S6: **Parameters obtained for thermodynamic and kinetic models.** Probability densities of the model parameters obtained via Bayesian statistics from two independent calculations for model1 (**a**), for model2 (**b**), and for model3 (**c**). The two shades of magenta, green, and blue denote two independent calculations.

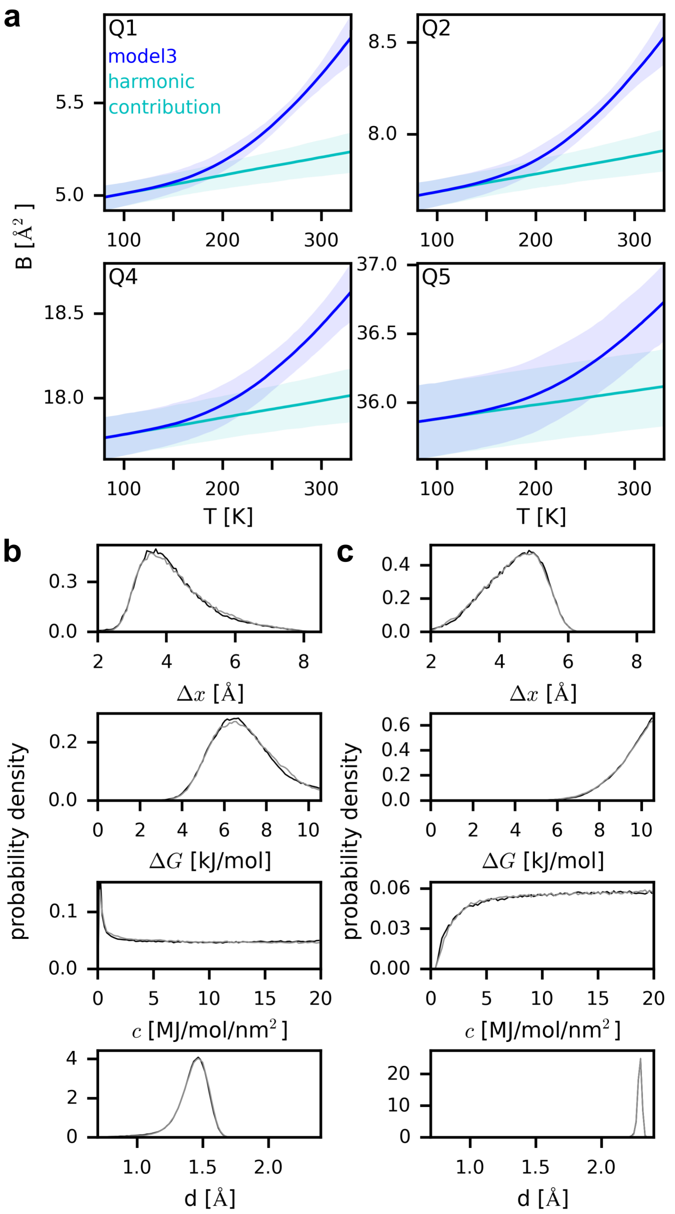


Figure S7: **B-factor dependency on temperature and model3 parameters.** (**a**) B-factors as a function of temperature calculated from model3 applied to ribosome T-quench simulations under equilibrium conditions (upper panel; expected values: blue lines, 95 % confidence interval: light blue area). The B-factors of the harmonic potentials are shown in cyan. (**b**,**c**) Probability densities of the parameters for the equilibrium model3 applied to experimental data. The B-factors obtained from x-ray crystallography of thaumatin[6] (b) and ribonuclease-A[7] (c) at different temperatures were used to train equilibrium model3.

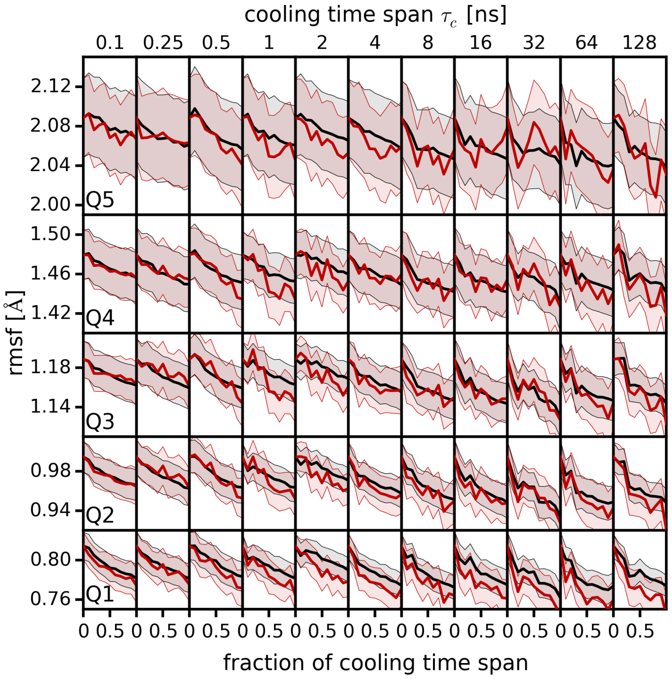


Figure S8: **Effect of additional sampling during T-quench simulations on rmsf distributions.** Rmsf quantiles are shown as a function the simulation time after rescaling (mean value: black line, standard deviation: gray area, compare Fig. 2c) and after an additional subtraction of the rmsf increase observed at T=277.15 K during the same simulation time (red line, light red area).

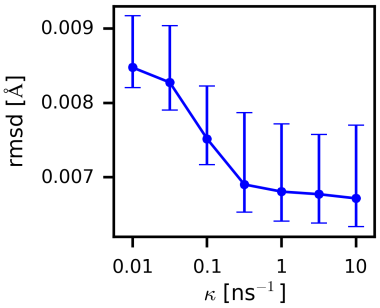


Figure S9: **Deviation of model3 rmsf values from the rmsf values obtained from T-quench simulations.** For different values of the scaling factor $\kappa$, the root mean square deviation (rmsd) of the predicted rmsf values from the rmsf values obtained from the T-quench simulations (Fig. 2c, black lines) is shown. The circles denote the mean rmsd values and the vertical lines correspond to the 95% confidence interval.

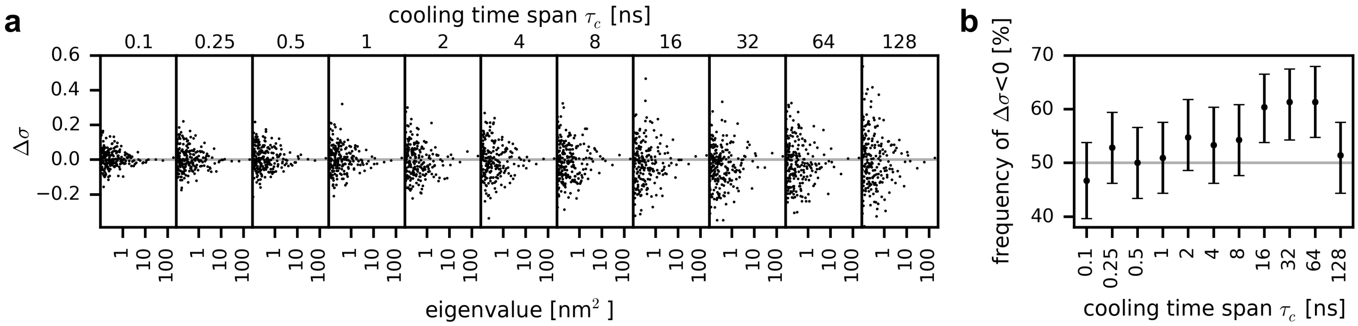


Figure S10: **Effect of cooling on conformational modes** (**a**) For each cooling time span, the reduction in the standard deviation $\Delta\sigma$ along conformational modes at the end of the T-quench simulation is shown. The conformational modes (eigenvectors) were obtained from a PCA of the trajectory at a temperature of 277.15 (Fig. S1a). For each eigenvector the eigenvalue and $\Delta\sigma = (\sigma_{\text{after}} - \sigma_{\text{before}})/\sigma_{\text{before}}$ are shown. $\sigma_{\text{before}}$ and $\sigma_{\text{after}}$ are standard deviations of the projections onto the eigenvector of the initial and final structures of the T-quench simulations, respectively (Fig. 2a, red and blue circles). (**b**) For each cooling time span, the percentage of eigenvectors for which $\Delta\sigma$ was below zero is shown. Bars denote the confidence intervals (95%) obtained from bootstrapping of the eigenvectors.
